# Multimodal cell type correspondence by intersectional mFISH in intact tissues

**DOI:** 10.1101/525451

**Authors:** Philip R. Nicovich, Michael J. Taormina, Christopher A. Baker, Thuc Nghi Nguyen, Elliot R. Thomsen, Emma Garren, Brian Long, Melissa Gorham, Jeremy A. Miller, Travis Hage, Alice Bosma-Moody, Gabe J. Murphy, Boaz P. Levi, Jennie L. Close, Bosiljka Tasic, Ed S. Lein, Hongkui Zeng

**Affiliations:** Allen Institute for Brain Science, Seattle, Washington

## Abstract

Defining a complete set of cell types within the cortex requires reconciling disparate results achieved through diverging methodologies. To address this correspondence problem, multiple methodologies must be applied to the same cells across multiple single-cell experiments. Here we present a new approach applying spatial transcriptomics using multiplexed fluorescence *in situ* hybridization, (mFISH) to brain tissue previously interrogated through two photon optogenetic mapping of synaptic connectivity. This approach can resolve the anatomical, transcriptomic, connectomic, electrophysiological, and morphological characteristics of single cells within the mouse cortex.

## Main

The parable of the Blind Men and the Elephant tells the story of a group of blind men, naïve to the nature of an elephant, presented with such a creature to try to conceptualize while observing only through touch (Figure 1A, upper). The men are each concerned with a single area and draw a conclusion for the whole based on their limited perspective. As scientists, we are similarly a group of observers each indirectly observing one portion of a large, hidden, and complex problem. The parable’s moral is that these challenging concepts may have many facets individually revealed to those with diverging perspectives. To create a comprehensive description, we must be able to synthesize these unique perspectives into a consistent whole.

**Figure 1.**
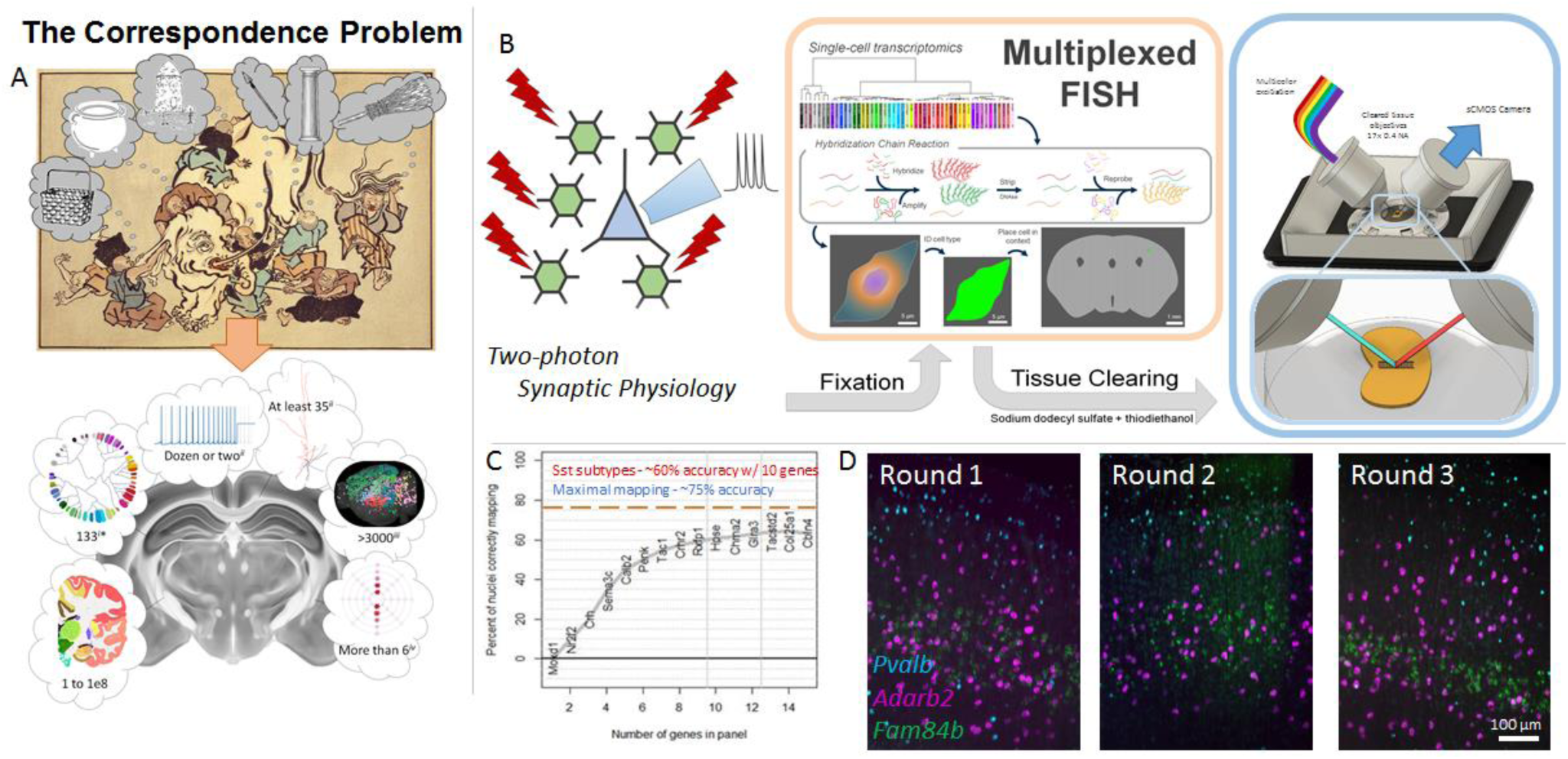
Overview of mFISH Light Sheet pipeline. A – The Correspondence Problem, demonstrated by the parable of the Blind Men and the Elephant. A modern scientific manifestation of this problem is generating agreement between cell type number, transcriptomic identity, physiology, morphology, connectivity, and function in the cortex. B – mFISH Light Sheet workflow. Tissue from an upstream modality is fixed, stained for mFISH, and imaged on an iSPIM microscope. Multiple color channels and rounds of stripping and reprobing against new transcript targets builds up a dataset of transcript abundance for target cells in the light sheet volumes. C – Bootstrap analysis to estimate mapping success as a function of gene information input in Sst cell types. Asymptote at ∼60% correct mapping to transcriptomic leaf nodes is achieved in 9 genes, versus 75% correct mapping with ‘all’ (thousands of) genes. D – Repeated probing of a 350 µm thick slice by HCR and imaged by light sheet microscopy. All genes are visible in expected patterning across all rounds.

This challenge of differing perspectives leading to seemingly conflicting outcomes is exemplified in attempts to understand the number and features of neuronal cell types within the mouse cortex (Fig 1A, lower). Recent studies report self-consistent outcomes based on anatomy^1^, transcriptomics^2,3^, electrophysiology^4^, morphology^4^, long-range^5^ and local connectomics^6^, and *in vivo* recordings^7^. However, these studies markedly disagree on the number of underlying cell types. Even if they did agree on the number of types, how results from one study map to those of another is not obvious. To reconcile this problem of cell type correspondence, single-cell characterization must traverse multiple modalities.

Each of these modalities, save transcriptomics, explicitly or implicitly carries spatial information within the specimen at single-cell resolution. Multiplexed fluorescence *in situ* hybridization (mFISH) enables a transcriptomic readout with this same single-cell spatial resolution. To link transcriptomic profiling with other upstream experimental modalities, we have extended the hairpin chain reaction (HCR) approach^8^ with DNase digestion for up to three rounds of hybridization against target transcripts with three spectrally-defined probes in each round. This protocol is compatible with paraformaldehyde-fixed mouse slices up to 350 µm thick, including those having undergone previous experiments.

To ensure cells interrogated in upstream methods can be tracked across subsequent experiments we chose to focus on those microscopy methods for mFISH readout that can be performed in 350 µm thick tissue samples without resectioning. For adult mouse brain tissue of this thickness, tissue clearing is required to reduce the large amount of scattering in native tissue, which is especially problematic in single-photon microscopy. Clearing is readily achieved through incubation in index-matching fluid comprised of 67% thiodiethanol (TDE) in PBS. An inverted selective plane illumination microscope (iSPIM)^9,10^ equipped with long-working-distance objectives compatible with the geometry and immersion fluid requirements (Special Optics, 16.7x/0.4 NA) yields rapid, full-thickness images at sub-cellular resolution and high speed. Multicolor, full-thickness image volumes over many square millimeters of lateral area are captured in under an hour, an order of magnitude faster than achievable with a point-scanning confocal at similar resolution.

A custom computational pipeline efficiently transforms these large, multi-color and multi-position 3D datasets from the as-acquired image stacks into aligned and stitched volumes readily explored through a web browser interface^11^. With each round of mFISH imaging as a separate acquisition volume, target cell locations from the upstream modality can map to the same cells in each volume. Extracting fluorescent signals across all channels and rounds yields a transcriptomic profile for each target cell.

These advancements allow the spatial transcriptomics portion of the pipeline to intersect with a range of upstream modalities. These modalities must meet a set of the following compatibility criteria: 1) the upstream modality must yield a tissue specimen compatible with downstream mFISH processing and imaging; 2) target cells of interest must be tagged with a fluorescent marker or markers that can be read out in both the upstream and mFISH modalities; 3) these target cells each must have a recorded spatial location within the specimen; and 4) these target cells should be sufficiently sparse for reliable mapping of upstream target cell locations into the mFISH volume coordinate space.

Two photon optogenetic mapping of synaptic connectivity provides an exemplar use case for this approach^12–15^ (Fig 1B). This modality uses sensitive channelrhodopsins amenable to two photon stimulation, such as ReaChR^16^ or Chrimson^17^, to probe cells for potential pre-synaptic connections to a selected patch-clamped cell. Dozens or hundreds of channelrhodopsin-expressing pre-synaptic cells can be probed in rapid succession for increased throughput compared to paired recordings. These experiments are performed in acute 350 µm-thick brain slices from animals expressing the channelrhodopsin and reporter fluorescent protein in a target cell subclass, and the patched cell passively filled with AlexaFluor dye and biotin through the duration of the patch recording. Post-experiment slices are fixed in paraformaldehyde in anticipation of further processing.

As a demonstration of our approach, we focus on cells marked by somatostatin (*Sst*), a subclass of inhibitory neurons. Channelrhodopsin is expressed in a Cre-dependent manner, either by delivery of a virus or breeding of a reporter mouse to *Sst-IRES-Cre* mice ^18,19^. This provides a good deal of *a priori* information about the transcriptomic identity of the target cells (as well as ensuring the sparsity of target cells desired for both the mFISH mapping and two photon optical stimulation steps). However, *Sst* is a broad marker with 21 identified transcriptomic types in mouse cortex^2,3^. Post-hoc transcriptomic refinement with mFISH against selected transcriptomic targets can disambiguate the cell types within this subclass observed in a specimen.

A bootstrap approach to probe target selection yields an ordered list of marker transcripts best able to discern the different *Sst* types (see mFISH gene selection in Methods). Given a reference single-cell RNA sequencing dataset^2^, mFISH-specific filters (including range of expression in expressing and non-expressing cells, mRNA length, and differential expression between types) refine the possible transcript target pool^20^. Selection begins with manually-selected marker genes (here *Sst* only) and additional targets from the filtered pool iteratively added while testing the confidence of single cells which include only selected gene targets mapping to the taxonomy built with the full reference dataset. The gene which most increases this mapping confidence is included in the selected target pool and the process repeats for choosing the next target.

This bootstrap method yields an asymptotic approach to approximately 60-65% correct mapping to an *Sst* sub-type using information only from *Sst* (provided by the optogenetic reporter) and 9 other genes. It is noteworthy that with the “full” transcriptomic data set single cells map to the correct type approximately 75% of the time. Those incorrect mappings are nearly always due to inconclusive mappings at one node from the type terminus or to a neighboring, closely-related type. With this it is apparent that three rounds of mFISH with three transcripts probed each round should be sufficient for disambiguation within a cell subclass, even within one as broad as the *Sst*-expressing interneurons.

In a wild-type mouse specimen we probed three transcripts repeatedly across three rounds (Fig 1D). These transcripts – *Pvalb, Adarb2*, and *Fam84b* – were chosen as genes with example ‘high’, ‘medium’, and ‘low’ expression levels and with characteristic expression patterns (*Pvalb* and *Adarb2* in mutually-exclusive lineages of inhibitory neurons and *Fam84b* in cortical layer 5 excitatory neurons)^2,3,20^. Fluorophores and HCR initiator sequences corresponding to a given target were cycled across rounds such that incomplete stripping between rounds or failed re-probing would be apparent as errant signal. In each round, the specimens were imaged on the iSPIM microscope and data passed through the data processing pipeline. The resulting images show that across three rounds all three probes are successfully repeatedly probed. The pattern of expression for each target is as expected in all three rounds. Given that the lowest-expressed gene, *Fam84b*, gives strong signal even in the final round, there is minimal apparent mRNA loss even after two DNase stripping cycles and 13 days of mFISH processing and imaging.

Intact specimens carried forward from two photon mapping of synaptic connectivity experiments are similarly successfully probed through mFISH. We performed a single-round experiment in a brain slice from an *Sst-IRES-Cre;Ai136* mouse, with the filled cell labeled with AlexaFluor 488 dye and enhancement of the fill by labeling the biotin with streptavidin-AlexaFluor 488 (Figure 2A). Capturing this image volume of ∼3 mm × 2 mm × 0.5 mm with sampling 370 nm × 740 nm × 260 nm across 5 spectral channels was completed within 45 minutes. The patch clamp recordings and morphology of the filled cell indicate that this is an excitatory neuron. HCR signal in this cell confirms expression of *Slc17a7*, a canonical marker for excitatory cells. Within this specimen, all cells with *Sst* HCR signal also show signal for *Gad2*, a pan-inhibitory marker. As expected, we observe that many *Gad2-*positive cells are *Sst-*negative, consistent with the expectation that *Gad2* labels all inhibitory neurons, of which *Sst*-expressing cells are a subset. In this specimen, cells labeled by the optogenetic *Sst-IRES-Cre*-dependent ReaChR-EYFP reporter are a further subset of cells showing *Sst* expression by mFISH.

**Figure 2.**
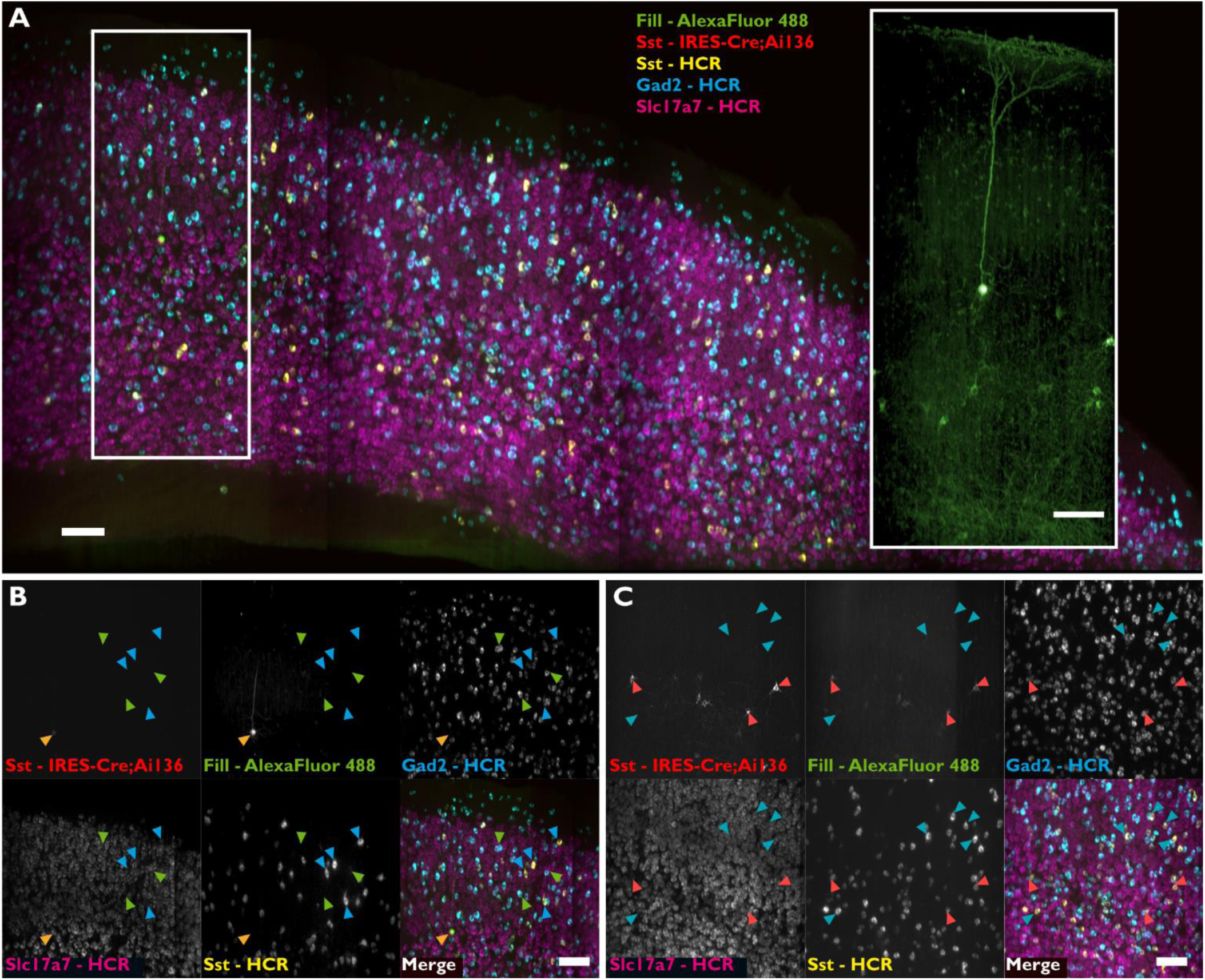
Cell class identification by mFISH Light Sheet. A – Maximum intensity projection of a 3 mm × 2 mm × 0.35 mm region of a cleared mouse brain sagittal slice imaged on the light sheet microscope. Signal for a filled cell from upstream a synaptic physiology experiment, Cre-dependent opsin reporter, and three transcripts probed by HCR is visible. Inset shows box area in fill and reporter channel only. B – Filled cell (orange marker) shows expression of Slc17a7 and not Gad2, confirming excitatory cell morphology observed in A, inset. All Sst-positive cells (blue) show expression of Gad2, but not all Gad2-positive cells show Sst expression (Gad+/Sst-, green markers). C – Opsin reporter-positive cell all show Sst expression (reporter+/Sst+ vermillion markers) but not all Sst-positive cells show opsin reporter signal (reporter-/Sst+ turquoise markers). Scale bars correspond to 100 µm.

A specimen expressing a nominally *Sst-IRES-Cre* dependent optogenetic reporter, ChrimsonR, shows an interesting contrast^18^ (Figure 3). Target cells within this specimen were probed through the synaptic connectivity mapping workflow and single-gene mFISH probing. Again, the filled cell (AlexaFluor 488 fill and streptavidin-AlexaFluor 488) and soma-localized opsin reporter (Chrimson-EYFP) were visible in the mFISH image volume. Our computational workflow maps the synaptic physiology target cell coordinate set to cells within the mFISH volume. Extracted sub-volumes around each target cell yield cell boundaries within the light sheet image volume based on the reporter signal channel (Fig 3C). This mask applied to the HCR channel yields information about the gene expression in the target cells.

**Figure 3.**
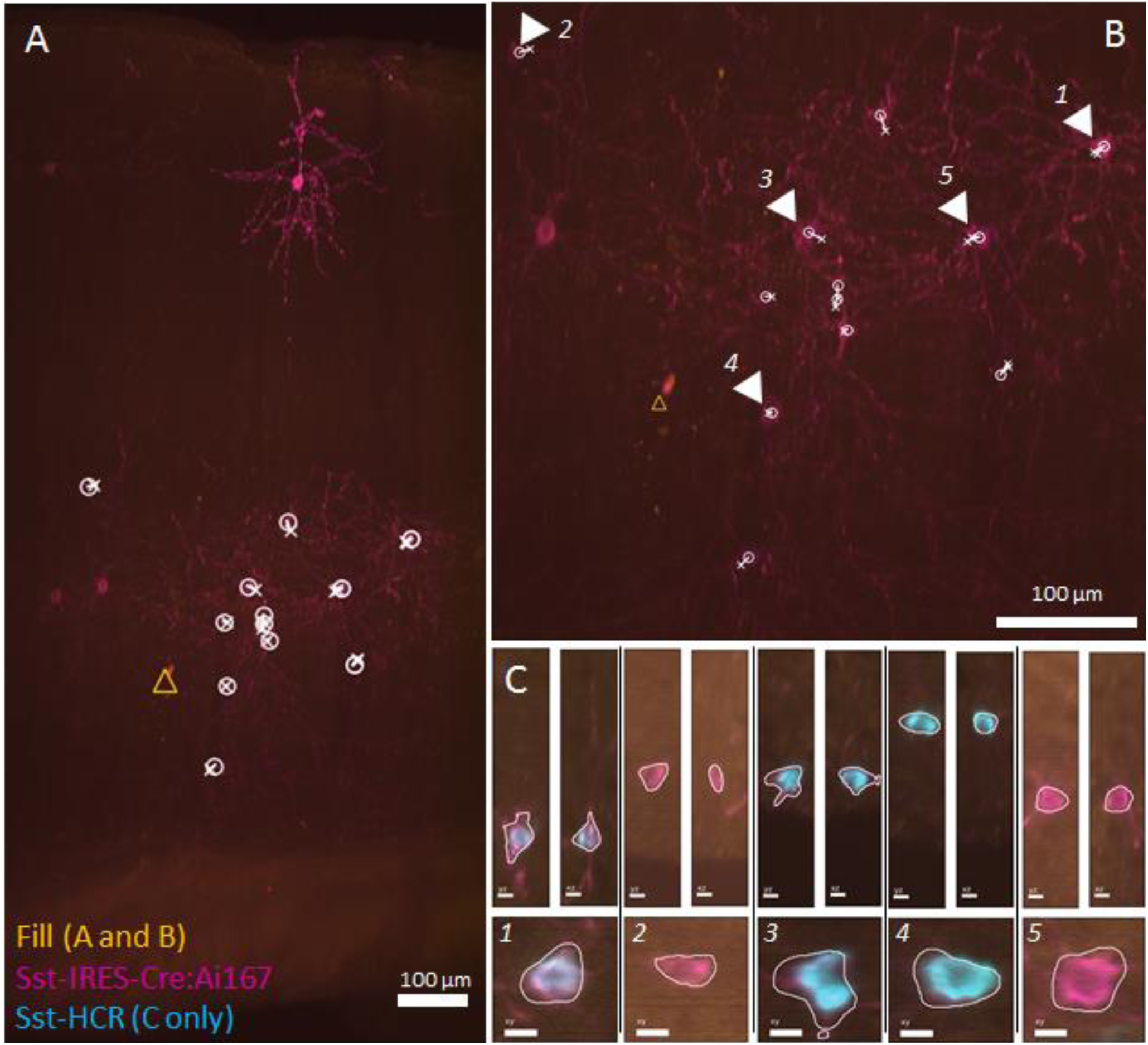
mFISH Light Sheet on a Two photon Synaptic Physiology slice. A – Slice previously probed by Synaptic Physiology, stained for Sst by HCR (shown in C only), and imaged on the iSPIM microscope. Full-thickness maximum intensity projection of 350 µm thick slice shown. Patched and filled cell indicated by yellow circle. White ‘X’ indicates manual alignment of upstream target point cloud; white circles indicate computer-assisted alignment; yellow triangle marks filled cell location. B – Inset showing region of slice with patched cells. C – ROIs taken around indicated target cells. White outlines indicate automatic segmentation boundary with plane shown corresponding to center of mass of segmented region. Note not all Sst-Cre reporter positive cells contain signal for Sst HCR.

In this specimen, a cell with opsin reporter signal does not always carry observable mFISH signal for *Sst*. This difference could imply transitional or development-dependent Cre expression or other cell state *versus* cell type distinctions. This difference would not be clear without profiling both reporter and transcript signals through a method such as the one shown here.

The ultimate goal for this approach is to enable transcriptomic cell type identification of specific cells with clear connectomic results. We demonstrate an example of this desired outcome by first injecting a virus encoding Cre-dependent ChrimsonR-EYFP-Kv2.1^21,22^ into the brain of an *Sst-IRES-Cre* mouse. The channelrhodopsin and reporter is sparsely expressed in *Sst*-expressing cells. A brain slice from this specimen is analyzed using two photon optogenetic connectivity mapping, showing a connection between an optically-targeted cell (Fig 4A and 4B, red) and a patched cell. Neighboring cells show no detected connection (Fig 4A and B, blue and yellow) to the patched cell. These cells can be located across multiple rounds of HCR staining, light sheet imaging, and stripping (2^nd^ of 3 rounds shown in Fig 4 C and D). The HCR signal for each cell and color-round gives an expression profile across the 9 transcripts probed for each target cell.

**Figure 4.**
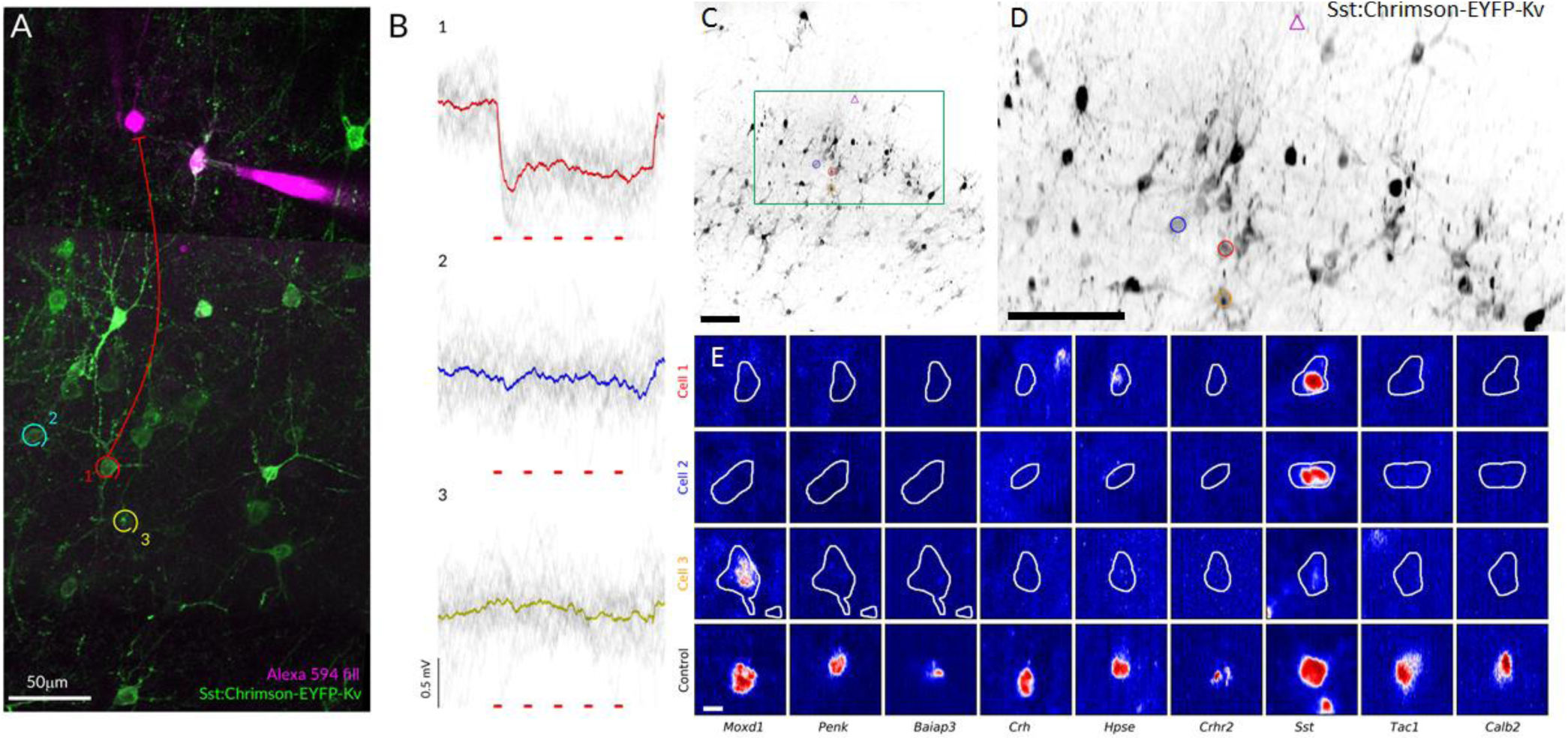
Investigating cell-type connection rates by mFISH Light Sheet – A – Image from two photon optogenetic connectivity mapping experiment. Pre-synaptic cells in question are marked in red (Cell 1), blue (Cell 2), and yellow (Cell 3) with Cell 1 demonstrating a connection to the indicated patched cell (magenta). B – Stimulation traces from connectivity mapping experiment for the three cells indicated in A. Cell 1 (red) shows a connection to patched cell. Red bars indicate 10 ms photostimulation epochs. C – Slice view of reporter fluorescence on light sheet microscope. D – Inset of C. Cells marked with circles correspond to A. Diamond is location of patched cell. E – Image-based heat map of log transformed intensity for all 9 genes across three profiled cells. White outline is border of cell defined by reporter fluorescence. Control row is a positive cell for each profiled gene. Each column colormap is normalized to positive control cell signal.

All nine probed transcripts show signal across this three-round experiment, confirming that the transcriptomic resolution required to map to the cell type level is achievable (Fig 1C), though further refinement may be required for the specific case of mapping *Sst* subtypes. The target cell showing a connection to the patched cell does show expression for *Sst* as well as moderate expression for *Hpse*, a marker for a transcriptomically-distinct *Sst* sub-type. The other two target cells do not share this same expression profile. Cell 2 does not show *Hpse* expression and Cell 3 has much lower *Sst* expression with clear *Moxd1* expression. These results show that the combination of two photon optogenetic connectivity mapping and mFISH can discern the differences in transcriptomic state of target cells within a broad interneuron subclass. With sufficient accumulation of connectomics data, enabled by the increased throughput of two photon optogenetic connectivity mapping, can yield cell-type-specific rates for local neuronal connectivity.

Potentially this approach can be generalized to intersect with other modalities operating on a single piece of tissue, given the specific constraints for downstream mFISH-based transcriptomic profiling. Especially beneficial is the rapid light sheet imaging step which, in principle, can cover an unlimited lateral area and up to 4 mm deep, making this a prime approach for *in situ* screening of viral or transgenic tools in conjunction with cell-type-level transcriptomic profiling of target cells Even without an upstream intersecting modality the demonstrated mFISH approach is an effective means to discern this information. Including an intersecting modality can reveal how subtle cell type differences correspond to differences in connectomics, morphology, function, or other metrics within the brain.

## Acknowledgements

We thank the Allen Institute founder, Paul G. Allen, for his vision, encouragement and support. We thank Jon Daniels and Applied Scientific Instrumentation, Inc., Harry Choi and Molecular Technologies, and Robert Campbell and Michael Davidson for support of instrumentation and reagents for this work. The authors acknowledge program management support of Dr. Susan Sunkin to enable these studies.

## Methods

### Animals and transgene expression

All animal housing and procedures were conducted under protocols consistent with the National Institutes of Health guidelines and approved by the Institutional Animal Care and Use Committee at the Allen Institute (Seattle, WA). To establish the general utility of our mFISH pipelines we examined animals from multiple sources, including Sst-IRES-Cre driver mice (JAX Stock # 013044) crossbred with ROSA26-ZtTA/J mice^23^ (JAX Stock # 012266) and Ai136 mice^18^, in which a fusion of the ReaChR opsin^16^ with EYFP is expressed from the TIGRE locus^19,24^ in a Cre- and tetracycline transactivator (tTA)-dependent manner (Figure 2). We also examined Sst-IRES-Cre driver mice crossed with the new Ai167 line with both the ChrimsonR opsin^17^ and the tTA contained within the new TIGRE 2.0 cassette^18^ (Figure 3 experiments). In other experiments (Figure 4), ∼P28 Sst-IRES-Cre driver mice were anesthetized with an isoflurane/oxygen mixture and 200nL of adeno associated virus (AAV) serotype 1 (∼5×10^12^ genome copy units/mL) was injected through a small craniotomy at 0.3 mm and 0.6 mm beneath the surface of primary visual cortex (3.16mm anterior and 2.75mm lateral to bregma). The AAV delivered a bicistronic construct consisting of ChrimsonR fused to a Kv2.1 somatic localization motif ^21,22^ that has been shown to enhance resolution and sensitivity for two photon optogenetic mapping experiments ^15^. Nuclei of opsin-positive cells were visualized using a Histone2B fusion protein with mTFP1 (a gift from Robert Campbell & Michael Davidson; Addgene plasmid # 54553; http://n2t.net/addgene:54553;RRID:Addgene_54553; Ai at al., 2006).

### Tissue preparation for synaptic physiology

Recording preparations were made per established pipelines for electrophysiological characterization of mouse neurons^4,6^, and all solutions were maintained under 95% O2/ 5% CO2. Under deep isoflurane anesthesia, mice (P40-60) of either sex were transcardially perfused with cold artificial cerebrospinal fluid (aCSF-1) containing (in mM): 98 HCl, 96 N-methyl-d-glucamine (NMDG), 2.5 KCl, 25 D-Glucose, 25 NaHCO3, 17.5 4-(2-hydroxyethyl)-1-piperazineethanesulfonic acid (HEPES), 12 N-acetylcysteine, 10 MgSO4, 5 Na-L-Ascorbate, 3 Myo-inositol, 3 Na Pyruvate, 2 Thiourea, 1.25 NaH2PO4, 0.5 CaCl2, and 0.01 taurine. Parasagittal slices 350 µm thick were prepared in cold aCSF-1 with a Compresstome (Precisionary Instruments) using a 17° slicing angle to preserve apical dendrites of neurons in primary visual cortex. Slices were maintained in a holding chamber (BSK 12, Scientific Systems Design) containing oxygenated aCSF-1 at 34°C for 10 min (Ting et al., 2014; Hájos and Mody, 2009) before transfer to ambient temperature holding solution (aCSF-2) containing (in mM) 94 NaCl, 25 D-Glucose, 25 NaHCO3, 14 HEPES, 12.3 N-acetylcysteine, 5 Na-L-Ascorbate, 3 Myo-inositol, 3 Na Pyruvate, 2.5 KCl, 2 CaCl2, 2 MgSO4, 2 Thiourea, 1.25 NaH2PO4, 0.01 Taurine for a minimum of one hour prior to recording.

### Electrophysiology and optogenetic mapping

Electrophysiological recordings were conducted at 31–33°C in custom chambers perfused (2 mL/min) with aCSF-5 containing (in mM): 2 CaCl2, 12.5 D-Glucose, 2 MgSO4, 1.25 NaH2PO4, 3 KCl, 18 NaHCO3, 126 NaCl, 0.16 Na L-Ascorbate. Filamented borosilicate glass pipettes were pulled to a tip resistance of 3–8 MΩ (diameter ∼ 1.25 µm), and filled with internal solution containing (in mM): 130 K-gluconate, 10 HEPES, 0.3 ethylene glycol-bis(β-aminoethyl ether)-N,N,N’,N’-tetraacetic acid (EGTA), 3 KCl, 0.23 Na2GTP, 6.35 Na2Phosphocreatine, 3.4 Mg-ATP, 13.4 Biocytin, and either 50 µM Alexa-594 or Alexa-488 hydrazide (Thermo Fisher Scientific). Signals were amplified using Multiclamp 700B amplifiers (Molecular Devices) and digitized at 50–200 kHz using ITC-1600 digitizers (Heka). Custom software, Multi-channel Igor Electrophysiology Suite (MIES; https://github.com/AllenInstitute/MIES), written in Igor Pro (WaveMetrics), was used for data acquisition.

For two photon mapping experiments, 1-4 neurons were patched and their intrinsic properties measured at their normal resting potential before maintaining cells at either −70mV or −55mV *via* current injection to isolate excitatory or inhibitory postsynaptic potentials, respectively. Virally transduced cells were visualized through a 40× 1.0 NA water immersion objective (Zeiss) on a two photon laser scanning microscope (Bruker Corp) with a tunable pulsed Ti:Sapphire laser (Chameleon Ultra, Coherent). Desired cells were individually photostimulated with a 1060 nm pulsed laser (Fidelity Femtosecond, Coherent) using galvo mirrors controlled by PrairieView software (Bruker Corp) to steer the beam through five revolutions of a 10 μm in diameter spiral over a 10 ms duration. Stimulation power was controlled by a Pockels cell (Conoptics) and was maintained at a level that elicited action potentials in each of 10 trials across > 90% of the cells tested. This varied as a function of opsin and transgene delivery but ranged from 18-85mW at the objective. Control experiments established that displacement of the stimulation focus from a recorded opsin-positive neuron by 12 µm laterally or 27 µm axially resulted in a >50% decrease in probability of eliciting an action potential, consistent with the resolution reported by other investigators^12,14^. Stimulation of putative presynaptic cells was repeated 20 times to test the stability of each putative connection and allow averaging to identify small amplitude potentials. Connections were scored by manual annotation using a voltage deconvolution technique and signal to noise threshold as previously described^6^.

### mFISH gene selection

Gene panels are selected using a combination of manual and algorithmic based strategies, and require a reference single cell/nucleus RNA-seq data set from the same tissue (in this case, ∼12,000 single cells in mouse primary visual cortex^2^). First, an initial set of high-confidence marker genes are selected through a combination of literature search and analysis of the reference data. These genes are used as input for a greedy algorithm (detailed below). Second, the reference RNA-seq data set is filtered to only include genes compatible with mFISH. Retained genes need to be 1) long enough to allow probe design (> 960 base pairs); 2) expressed highly enough to be detected (FPKM >= 10), but not so high as to overcrowd the signal of other genes in a cell (FPKM < 500); 3) expressed with low expression in off-target cells (FPKM < 50 in non-neuronal cells); and 4) differentially expressed between cell types (top 500 remaining genes by marker score^20^). To more evenly sample each cell type, the reference data set is also filtered to include a maximum of 50 cells per cluster.

The main step of gene selection uses a greedy algorithm to iteratively add genes to the initial set. To do this, each cell in the filtered reference data set is mapped to a cell type by taking the Pearson correlation of its expression levels with each cluster median using the initial gene set of size n, and the cluster corresponding to the maximum value is defined as the “mapped cluster”. The “mapping distance” is then defined as the average cluster distance between the mapped cluster and the originally assigned cluster for each cell. In this case a weighted cluster distance, defined as one minus the Pearson correlation between cluster medians calculated across all filtered genes, is used to penalize cases where cells are mapped to very different types, but an unweighted distance, defined as the fraction of cells that do not mapped to their assigned cluster, could also be used. This mapping step is repeated for every possible n+1 gene set in the filtered reference data set, and the set with minimum cluster distance is retained as the new gene set. These steps are repeated using the new get set (of size n+1) until a gene panel of the desired size is attained. Code for reproducing this gene selection strategy is available as part of the mfishtools R library (https://github.com/AllenInstitute/mfishtools).

### mFISH sample preparation

Slices were fixed in 4% PFA for 2 hours at room temperature (RT), washed three times in PBS for 10 min each, then transferred to 70% EtOH at 4°C for a minimum of 12 hours, and up to 30 days. Slices were then incubated in 8% SDS in PBS at RT for two hours with agitation. The solution was exchanged with 2X SSC three times, slices were washed for one hour at RT, followed by two additional one hour washes with fresh 2X SSC.

### In situ HCR

We performed HCR v3.0 using reagents and a modified protocol from Molecular Technologies and Molecular Instruments^8^. Slices were incubated in pre-warmed 30% probe hybridization buffer (30% formamide, 5X sodium chloride sodium citrate (SSC), 9 mM citric acid pH 6.0, 0.1% Tween 20, 50 µg/mL heparin, 1X Denhardt’s solution, 10% dextran sulfate) at 37°C for 5 min, then incubated overnight at 37°C in hybridization buffer with the first three pairs of probes added at a concentration of 4 nM (Table 1). The hybridization solution was exchanged 3 times with 30% probe wash buffer (30% formamide, 5X SSC, 9 mM citric acid pH 6.0, 0.1% Tween 20, 50 µg/mL heparin) and slices were washed for one hour at 37°C. Probe wash buffer was briefly exchanged with 2X SSC, then amplification buffer (5X SSC, 0.1% Tween 20, 10% dextran sulfate) for 5 min. Even and odd hairpins for each of the three genes were pooled and snap-cooled by heating to 95°C for 90 seconds then cooling to RT for 30 min. The hairpins were then added to amplification buffer at a final concentration of 60 nM, and slices were incubated in amplification solution for 4 hours at RT. This was followed by a brief wash with 2X SSC and a one hour, RT incubation in 2X SSC containing 8 µg/µl Brilliant Violet 421™ Streptavidin (BioLegend, Cat. No. 405225) and 0.05% Tween 20. Slices were washed three times for 10 min in 2X SSC.

For each round of imaging, an aliquot of 67% 2,2’-Thiodiethanol (TDE) solution was prepared for use as a clearing and immersion fluid. ≥99% TDE (Sigma-Aldrich) was mixed with DI water to create a 67% TDE solution with a refractive index of 1.46, verified by a pocket refractometer (PAL-RI, Atago). Slices were transferred to 67% TDE and allowed to equilibrate for at least 1 hour at room temperature prior to imaging.

### iSPIM imaging

Slices were mounted by transferring to a 25 mm circular, refractive index-matching (RI = 1.46) quartz coverslip (01019-AB, SPI Supplies) secured into a custom 3D printed coverslip holder (reference). The coverslip was then placed on a 25 mm round fused silica mirror blank (PF10-03, Thorlabs), placed in the center of a custom 3D printed large volume sample chamber (https://github.com/PRNicovich/3D-Printed-Optics-Lab-Parts/tree/master/ASI%20Large%20Volume%20Chamber), such that the sample was positioned in between the coverslip and the mirror blank. A syringe was used to fill any air pockets with 67% TDE solution. The coverslip holder containing the coverslip was loosely screwed in to the bottom of the sample chamber to secure the sample.

Imaging was performed using an Inverted Selective Plane Illumination Microscope (iSPIM) by Applied Scientific Instrumentation (ASI) configured as a single-sided system and equipped with an XY automated stage (MS-2000, ASI). The sample chamber, in place of a stage insert, was mounted on the stage and filled with 67% TDE solution. Microscope controls and image acquisition were carried out using custom in-house scripts and the ASI diSPIM Device Control plugin for Micro-Manager 1.4.

Multicolor stacks were acquired using a stage scan acquisition mode and a slice step size of 0.748µm. At this spacing, pixels from neighboring frames in the stack are aligned vertically (perpendicular to the slice plane) with a modulus of four pixels. An illumination objective (Cleared Tissue Objective, 16.7x, 0.4 NA, 12mm WD, ASI) was used to generate a light sheet. Brilliant Violet 421 was excited with a Vortran Stradus-405-100 laser and Semrock FF02-447/60 emission filter. mTFP1 and Alexa-488 were excited with a Stradus-488-150 laser (Vortran) and Olympus U-MWG emission filter (510-550 nm). EYFP was excited with a Stradus-514-60 laser (Vortran) and Semrock FF01-550/32 emission filter. Alexa-546 was excited with a Stradus-561-50 laser (Vortran) and Semrock FF01-580/23 emission filter. Alexa-594 was excited with a MGL-III-589 50mW laser (Changchun New Industries) and Semrock FF01-615/24 emission filter. Alexa-640 was excited with a Stradus-642-110 laser (Vortran) and Semrock FF01-432/515/595/730 emission filter.

Laser powers were maintained at <10mW for all imaging experiments and modulated through a NicoLase laser sequencer^25^. Fluorescence was detected through a second, identical cleared tissue objective and imaged onto a digital CMOS camera (ORCA-Flash4.0, Hamamatsu) at 50 ms exposure time. Raw acquired data was immediately saved to a network attached storage (NAS) device as sequential concatenated TIFF stacks, split into files 4.2 GB in size.

### Stripping and subsequent hybridization rounds

To strip the signal in preparation for subsequent rounds, 67% TDE was exchanged with 2X SSC three times and samples were washed for 1 hour. 2X SSC was replaced with 1X DNase buffer for 5 min and then a 1:50 dilution of DNase I in DNase buffer (DNase I recombinant, RNase-free, Roche, Cat. No. 04716728001), and incubated for 1 hour at 37°C. This solution was replaced with fresh DNase solution before incubating slices overnight at 37°C. Slices were washed with 65% formamide in 2X SSC for one hour at 37°C, then 2X SSC for one hour at RT, before being transferred to 67% TDE for at least one hour. After imaging to confirm the signal was gone, the slices were washed in 2X SSC for one hour to remove TDE before proceeding to subsequent hybridization rounds, which followed the protocol described above, except omitting the incubation in streptavidin solution.

### Computational data processing pipeline

As-acquired TIFF stacks were each resliced into individual planes along the *xz* plane of the camera coordinate space, corresponding to planes parallel to the *x’y’* plane of the stage coordinate space. Each resliced plane is 2048 pixels by approximately 500 pixels in size and contains data for one color channel.

These planes must be deskewed from the oblique camera perspective into the stage coordinate space. The stage step size above is purposefully chosen such that cumulatively translating every second plane along the camera *z* axis by one pixel in the *y* axis closely approximates this skew transformation. This approach produces negligible errors on the scale of a few pixels due to the discrete shear, but is a more efficient calculation than, for example, an affine transform for datasets of this scale.

The *x’y’z’* position of each resliced plane in the image volume is recorded in a Render tilespec JSON file (https://github.com/saalfeldlab/render). This position includes the deskew transformation applied to the relevant planes. An updated transformation, such as to correct for stage alignment between imaged strips or color channels, applies to the tilespec JSON file and not the original planes. Given a specified bounding box in the image volume the Render server returns the desired voxels on-the-fly from discrete planes stored on disk.

Pre-computing a multi-resolution image pyramid files using Cloud-Volume and Igneous python libraries (https://github.com/seung-lab) aids in rapid visualization of the image volumes. For each data set we generate this pre-computed volume and can then view these through NDViz on a standard web browser^11^. Projection images along each axis and for each imaged color channel are generated through this pre-computed volume for use in the cross-modality alignment.

Targeting coordinates from the upstream synaptic physiology experiment metadata are transformed into coordinates in units of microns with positions relative to the filled cell. This coordinate point cloud is overlaid on an *x’y’* maximum intensity projection of the filled cell and reporter fluorescence channels from the light sheet microscope acquisitions. The coordinate point cloud rotates about the pinned filled cell position to find a maximal correspondence between the point cloud and filled cells in the reporter projection image. A corollary workflow finds updates the correspondence transform of the point cloud against *y’z’* and *x’z’* projections. Once a user finds these rough alignments, an automated non-linear distortion locates the final target cell positions in the imaging volume. This non-linear distortion finds the closest regional maximum within a bounding box (typically 25 µm × 25 µm around the rough point position) of the intensity of the reporter projection image following the application of a uniformity filter (scipy.ndimage.filters.uniform_filter; scale of 15 µm).

Sequential rounds of mFISH staining and light sheet imaging independently travel through this correspondence workflow. The final output is a set of point clouds where each point is associated with a target cell in the image space volume for a given round of light sheet imaging. The Render service returns a sub-volume (typically 50 µm × 50 µm × 150 µm) of all acquired color channels for each point an in each imaging round. A Frangi filter applied to the reporter channel signal (skimage.filter.frangi; scale range = (2, 20) pixels, betas = (1, 2)) followed by Otsu thresholding defines the boundary of the target cell within the image sub-volume. The intensity of the mFISH signals within this binarized volume provides inputs for the transcriptomic assignment.

An example cell positive for the mFISH probe is chosen for each transcript probed. The signal for each pulled target cell within this color channel and the control cell is individually background-subtracted and log transformed before all being normalized to the same colormap range as the control cell signal. These sets of images for each transcript are assembled into the intensity-based heat map shown in Figure 4E.

## Supplemental Information

**Table 1.** HCR probes used in this study

| Gene name | Accession | Initiator | # Probes Pairs | Fluorophore |
| --- | --- | --- | --- | --- |
| Moxd1 | NM_021509.5 | B3 | 20 | AlexaFluor 546 |
| Penk | NM_001348209.1 | B4 | 20 | AlexaFluor 594 |
| Baiap3 | NM_001163270.1 | B5 | 20 | AlexaFluor 647 |
| Crh | NM_205769.3 | B2 | 13 | AlexaFluor 546 |
| Hpse | NM_152803.4 | B3 | 20 | AlexaFluor 594 |
| Crhr2 | NM_001288620.1 | B1 | 20 | AlexaFluor 647 |
| Tac1 | NM_001311060.1 | B3 | 10 | AlexaFluor 594 |
| Sst | NM_009215.1 | B2 | 11 | AlexaFluor 546 |
| Calb2 | NM_007586.1 | B1 | 20 | AlexaFluor 647 |

## References

(1) Lein, E. S.; Hawrylycz, M. J.; Ao, N.; Ayres, M.; Bensinger, A.; Bernard, A.; Boe, A. F.; Boguski, M. S.; Brockway, K. S.; Byrnes, E. J.; et al. Genome-Wide Atlas of Gene Expression in the Adult Mouse Brain. Nature 2006, 445, 168.

(2) Tasic, B.; Yao, Z.; Graybuck, L. T.; Smith, K. A.; Nguyen, T. N.; Bertagnolli, D.; Goldy, J.; Garren, E.; Economo, M. N.; Viswanathan, S.; et al. Shared and Distinct Transcriptomic Cell Types across Neocortical Areas. Nature 2018, 563 (7729), 72–78.

(3) Tasic, B.; Menon, V.; Nguyen, T. N.; Kim, T. K.; Jarsky, T.; Yao, Z.; Levi, B.; Gray, L. T.; Sorensen, S. A.; Dolbeare, T.; et al. Adult Mouse Cortical Cell Taxonomy Revealed by Single Cell Transcriptomics. Nat. Neurosci. 2016, 19 (2), 335–346.

(4) Gouwens, N. W.; Berg, J.; Feng, D.; Sorensen, S. A.; Zeng, H.; Hawrylycz, M. J.; Koch, C.; Arkhipov, A. Systematic Generation of Biophysically Detailed Models for Diverse Cortical Neuron Types. Nat. Commun. 2018, 9 (1), 710.

(5) Harris, J. A.; Mihalas, S.; Hirokawa, K. E.; Whitesell, J. D.; Knox, J.; Bernard, A.; Bohn, P.; Caldejon, S.; Casal, L.; Cho, A.; et al. The Organization of Intracortical Connections by Layer and Cell Class in the Mouse Brain. bioRxiv 2018, 292961.

(6) Seeman, S. C.; Campagnola, L.; Davoudian, P. A.; Hoggarth, A.; Hage, T. A.; Bosma-Moody, A.; Baker, C. A.; Lee, J. H.; Mihalas, S.; Teeter, C.; et al. Sparse Recurrent Excitatory Connectivity in the Microcircuit of the Adult Mouse and Human Cortex. bioRxiv 2018, 292706.

(7) de Vries, S. E. J.; Lecoq, J.; Buice, M. A.; Groblewski, P. A.; Ocker, G. K.; Oliver, M.; Feng, D.; Cain, N.; Ledochowitsch, P.; Millman, D.; et al. A Large-Scale, Standardized Physiological Survey Reveals Higher Order Coding throughout the Mouse Visual Cortex. bioRxiv 2018, 359513.

(8) Choi, H. M. T.; Beck, V. A.; Pierce, N. A. Next-Generation in Situ Hybridization Chain Reaction: Higher Gain, Lower Cost, Greater Durability. ACS Nano 2014, 8 (5), 4284–4294.

(9) Kumar, A.; Wu, Y.; Christensen, R.; Chandris, P.; Gandler, W.; McCreedy, E.; Bokinsky, A.; Colon-Ramos, D. A.; Bao, Z.; McAuliffe, M.; et al. Dual-View Plane Illumination Microscopy for Rapid and Spatially Isotropic Imaging. Nat. Protoc. 2014, 9 (11), 2555–2573.

(10) Kumar, A.; Christensen, R.; Guo, M.; Chandris, P.; Duncan, W.; Wu, Y.; Santella, A.; Moyle, M.; Winter, P. W.; Colon-Ramos, D.; et al. Using Stage- and Slit-Scanning to Improve Contrast and Optical Sectioning in Dual-View Inverted Light Sheet Microscopy (diSPIM). Biol. Bull. 2016, 231 (1), 26–39.

(11) Vogelstein, J. T.; Perlman, E.; Falk, B.; Baden, A.; Gray Roncal, W.; Chandrashekhar, V.; Collman, F.; Seshamani, S.; Patsolic, J. L.; Lillaney, K.; et al. A Community-Developed Open-Source Computational Ecosystem for Big Neuro Data. Nat. Methods 2018, 15 (11), 846–847.

(12) Prakash, R.; Yizhar, O.; Grewe, B.; Ramakrishnan, C.; Wang, N.; Goshen, I.; Packer, A. M.; Peterka, D. S.; Yuste, R.; Schnitzer, M. J.; et al. Two photon Optogenetic Toolbox for Fast Inhibition, Excitation and Bistable Modulation. Nat. Methods 2012, 9 (12), 1171.

(13) Izquierdo-Serra, M.; Hirtz, J. J.; Shababo, B.; Yuste, R. Two photon Optogenetic Mapping of Excitatory Synaptic Connectivity and Strength. iScience 2018, 8, 15–28.

(14) Packer, A. M.; Peterka, D. S.; Hirtz, J. J.; Prakash, R.; Deisseroth, K.; Yuste, R. Two photon Optogenetics of Dendritic Spines and Neural Circuits. Nat. Methods 2012, 9 (12), 1202.

(15) Baker, C. A.; Elyada, Y. M.; Parra, A.; Bolton, M. M. Cellular Resolution Circuit Mapping with Temporal-Focused Excitation of Soma-Targeted Channelrhodopsin. Elife 2016, 5, e14193.

(16) Lin, J. Y.; Knutsen, P.; Muller, A.; Kleinfeld, D.; Tsien, R. Y. ReaChR: A Red-Shifted Variant of Channelrhodopsin Enables Deep Transcranial Optogenetic Excitation. Nat. Neurosci. 2013, 16 (10), 1499–1508.

(17) Klapoetke, N. C.; Murata, Y.; Kim, S.; Pulver, S. R.; Birdsey-Benson, A.; Cho, Y.; Morimoto, T. K.; Chuong, A. S.; Carpenter, E. J.; Tian, Z.; et al. Independent Optical Excitation of Distinct Neural Populations. Nat. Methods 2014, 11 (3), 338–346.

(18) Daigle, T. L.; Madisen, L.; Hage, T. A.; Valley, M. T.; Knoblich, U.; Larsen, R. S.; Takeno, M. M.; Huang, L.; Gu, H.; Larsen, R.; et al. A Suite of Transgenic Driver and Reporter Mouse Lines with Enhanced Brain-Cell-Type Targeting and Functionality. Cell 2018, 174 (2), 465–480.e22.

(19) Madisen, L.; Garner, A. R.; Shimaoka, D.; Chuong, A. S.; Klapoetke, N. C.; Li, L.; van der Bourg, A.; Niino, Y.; Egolf, L.; Monetti, C.; et al. Transgenic Mice for Intersectional Targeting of Neural Sensors and Effectors with High Specificity and Performance. Neuron 2015, 85 (5), 942–958.

(20) Hodge, R. D.; Bakken, T. E.; Miller, J. A.; Smith, K. A.; Barkan, E. R.; Graybuck, L. T.; Close, J. L.; Long, B.; Penn, O.; Yao, Z.; et al. Conserved Cell Types with Divergent Features between Human and Mouse Cortex. bioRxiv 2018, 384826.

(21) Zhang, Z.; Feng, J.; Wu, C.; Lu, Q.; Pan, Z.-H. Targeted Expression of Channelrhodopsin-2 to the Axon Initial Segment Alters the Temporal Firing Properties of Retinal Ganglion Cells. PLoS One 2015, 10 (11), e0142052.

(22) Lim, S. T.; Antonucci, D. E.; Scannevin, R. H.; Trimmer, J. S. A Novel Targeting Signal for Proximal Clustering of the Kv2.1 K+ Channel in Hippocampal Neurons. Neuron 2000, 25 (2), 385–397.

(23) Li, L.; Tasic, B.; Micheva, K. D.; Ivanov, V. M.; Spletter, M. L.; Smith, S. J.; Luo, L. Visualizing the Distribution of Synapses from Individual Neurons in the Mouse Brain. PLoS One 2010, 5 (7), e11503.

(24) Zeng, H.; Horie, K.; Madisen, L.; PLoS …, P. M. N. An Inducible and Reversible Mouse Genetic Rescue System. 2008.

(25) Nicovich, P. R.; Walsh, J.; Böcking, T.; Gaus, K. NicoLase – An Open-Source Diode Laser Combiner, Fiber Launch, and Sequencing Controller for Fluorescence Microscopy. PLoS One 2017, 12 (3).

